# Momilactone B inhibits Arabidopsis growth and development via disruption of ABA and auxin signaling

**DOI:** 10.1101/2020.09.30.320655

**Authors:** Jianxin Wu, Jun Long, Xianhui Lin, Zhenyi Chang, Scott R. Baerson, Chaohui Ding, Xiaoyan Wu, Zhiqiang Pan, Yuanyuan Song, Rensen Zeng

## Abstract

In competition for limited resources, many plants release allelochemicals to inhibit the growth of neighboring plants. Momilactone B (MB) is a major allelochemical produced by rice (*Oryza sativa*), however its mode of action is currently unknown. We used Arabidopsis (*Arabidopsis thaliana*) as a model system to evaluate potential mechanisms underlying the inhibitory effects of MB on seed germination, seedling establishment and root growth through the use of confocal microscopy and the examination of transcriptional responses in MB-treated seedlings. In response to MB treatment, transcript levels for genes encoding several key ABA biosynthetic enzymes and signaling components, including the transcription factor ABA-INSENSITIVE 4 (ABI4), were dramatically increased. Additionally, *ABA insensitive 4* (*abi4*) mutant seedlings exhibited reduced susceptibility to exogenously-provided MB. Although the transcript levels of *DELLA* genes, which negatively regulate GA signaling, were significantly increased upon MB exposure, exogenous GA application did not reverse the inhibitory effects of MB on Arabidopsis germination and seedling development. Moreover, a reduction in seedling root meristematic activity, associated with reduced expression of auxin biosynthetic genes and efflux transporters, and apparent lowered auxin content, was observed in MB-treated root tips. Exogenous auxin applications partially rescued the inhibitory effects of MB observed in root growth. Our results indicate that MB suppresses Arabidopsis seed germination and root growth primarily via disruption of ABA and auxin signaling. These findings underscore the crucial roles played by phytohormones in mediating responses to allelochemical exposure.

**One-sentence summary:** Momilactone B, the key allelochemical of rice, inhibits Arabidopsis growth and development via disruption of ABA and auxin signaling, suggesting the crucial roles of phytohormones in plant allelopathy

## Introduction

To gain an advantage in the competition for limited light, water and nutrient resources, certain plants species inhibit the growth of neighboring plants through the release of chemical compounds. This phenomenon is termed allelopathy, and the inhibitory compounds released are referred to as allelochemicals (Macias *et al.*, 2007; Inderjit *et al.*, 2011). Allelochemicals released within the soil environment can inhibit diverse processes such as seed germination, cell elongation, cell division, nutrient acquisition and photosynthesis, and are thought to profoundly influence plant community structure and evolution through the loss of susceptible species via chemical interference, and by imposing selective pressure favoring more tolerant species (Bais *et al.*, 2003; Macias *et al.*, 2007). Allelopathic crop species and their residues can also be used as cost-effective weed management tools for sustainable agriculture systems which reduce the requirement for synthetic herbicide applications (Jabran et al., 2015). An increased understanding of plant allelopathy will therefore not only improve our ability to devise effective approaches for limiting the spread of invasive weeds and preserving native plant stands, but could also lead to the development of more sustainable weed management practices.

Rice (*Oryza sativa* L.) represents one of the world’s most important food crops, therefore yield losses due to weed infestations in rice cropping systems result in substantial economic costs worldwide (Siddique & Ismail, 2013). Currently, the most well-established and effective weed control options available for rice involve synthetic herbicide spray applications, however the extensive use of synthetic herbicides and other chemical pesticides in agricultural systems poses significant risks to both the environment and human health (Neve *et al.*, 2009). Allelopathic crop varieties have therefore generated significant interest for the development of alternative, environmentally sound weed control methods, and in a study performed by Dilday and coworkers more than 5000 rice varieties were screened, resulting in the identification of 191 rice varieties exhibiting significant allelopathic activity (Dilday *et al.*, 1994). Significant efforts have also been made to introduce allelopathic traits into cultivated rice varieties and to identify all of the active allelochemicals produced by rice plants. The putative allelochemicals identified from rice to date include alkaloids, phenolics, flavonoids, glucosinolates and terpenoids (Kato-Noguchi *et al.*, 2002; Seal *et al.*, 2004a; Seal *et al.*, 2004b). Seal et al. (2004b) isolated and identified twenty-five compounds from the root exudates of both allelopathic and non-allelopathic rice varieties. Interestingly, the contents of five phenolics, including caffeic acid, *p*-hydroxybenzoic acid, vanillic acid, syringic acid, and *p*-coumaric acid, were determined to be significantly higher in allelopathic rice varieties, however, the concentrations of these putative allelochemicals in rice root exudates and soils were much lower than what is thought to be required for the effective growth inhibition of weeds (Seal *et al.*, 2004a; Seal *et al.*, 2004b).

Momilactones A (MA) and B (MB) are highly active diterpene allelochemicals found in rice root exudates as well as hulls, and are among the most extensively studied (Kato et al., 1973; Takahashi et al., 1976; Kato-Noguchi *et al.*, 2002, 2008). Both compounds exert strong inhibitory effects on the growth of susceptible species, although MB exhibits significantly higher activity than MA (Kato-Noguchi *et al.*, 2010). For example, the IC_50_ values (concentration required for 50% growth inhibition) determined for barnyard grass (*Echinochloa crus-galli*) seedling shoot and root tissues were 146 and 91 μM for MA, and 6.5 and 6.9 μM for MB, respectively (Kato-Noguchi *et al.*, 2010). Although the overall content of MB in rice plants is less than that of MA, higher levels of MB are exuded by rice seedling root systems. Taking the relative activities of the two compounds into consideration, it has been estimated that MA accounts for only 0.8–2.2% of the observed growth inhibition of *E. crus-galli* by rice, whereas MB accounts for 59–82% of the observed growth inhibition, suggesting a major role for MB in the allelopathic potential of rice plants (Kato-Noguchi *et al.*, 2010). Furthermore, RNAi-mediated inhibition of key momilactone biosynthetic genes such as copalyl diphosphate synthase 4 (*OsCPS4*) and kaurene synthase-like 4 (*OsKSL4*), resulted in significant reductions in momilactone release and reduced allelopathic activity of rice seedlings in *in vitro* assays, and similarly, over-expression of *OsCPS2* and *OsCPS4* significantly increased allelopathic potential *in vitro* (Xu *et al.*, 2012; Niu et al., 2017). Taken together, these studies clearly demonstrate the key role played by momilactones in rice allelopathy, and particularly that of MB.

Although numerous studies have been conducted to date which document the physiological effects of momilactones on susceptible plant species, and/or address the ecophysiological roles they may play, a relative paucity of information exists concerning the mode of action of these compounds. In one study performed by Kato-Noguchi and coworkers (2013) examining the effects of MB on germinating Arabidopsis seedlings, a reduction of seed storage protein metabolism was observed. In their study, protein levels of subtilisin-like serine protease, amyrin synthase LUP2, β-glucosidase and malate synthase were significantly decreased, while those of glutathione S-transferase and 1-cysteine peroxiredoxin 1 were increased, indicating that MB may affect Arabidopsis early seedling growth by inhibiting the mobilization of protein storage reserves (Kato-Noguchi *et al.*, 2013). Direct evidence linking MB to a specific cellular target or targets however is still lacking.

Given the limited genetic information available for *E. crus-galli* and other noxious weeds commonly associated with rice cultivation, we instead employed Arabidopsis as a model to further investigate the *in vivo* mechanism of MB action. In the present work, the effects of MB exposure on seed germination, seedling establishment and root development were evaluated, and the mechanism of MB-induced inhibition of these processes was investigated. Our results indicated that MB exposure inhibits seed germination in Arabidopsis primarily through the induction of both the biosynthesis of ABA as well as ABA-mediated signaling components. Consistent with this, ABA– insensitive *abi4* mutant seeds were observed to exhibit dramatically higher germination frequencies in the presence of MB relative to wild-type seeds. Additionally, MB appeared to inhibit seedling root system development via the disruption of auxin biosynthesis and the polar transport of auxins within root tips. Application of exogenous auxin was observed to partially reverse the inhibitory effects of MB on roots. Collectively, our findings highlight the critical roles played by the phytohormones ABA and auxin in MB-mediated allelopathic inhibition.

## Results

### MB inhibits germination and early seedling establishment

The potential inhibitory effects of MB on Arabidopsis were first evaluated by comparing germination frequencies and cotyledon greening in the presence and absence of exogenously supplied MB (Fig. 1a-c). In the presence of 4 μM MB, the average % germination observed was approximately 69% at 36 hours post-planting, whereas approximately 99% of the seeds place on MB-free medium had germinated by the 36 h timepoint (Fig. 1a-b). Cotyledon greening, which represents a critical step for seedling establishment (Shu *et al.*, 2013), was also compared at 4 days post-planting for seedlings germinated in the presence or absence of MB. As was observed for germination frequencies, cotyledon greening was also significantly impaired by exposure to MB (Fig. 1a, c). Approximately 99% of the seedlings grown on MB-free medium had greened within the 4 day period, whereas seedlings grown on MB-supplemented medium exhibited no discernible greening within this time period. Thus, the results clearly demonstrate the dramatic inhibitory effects imposed by MB on Arabidopsis germination and early seedling establishment.

**Figure 1.**
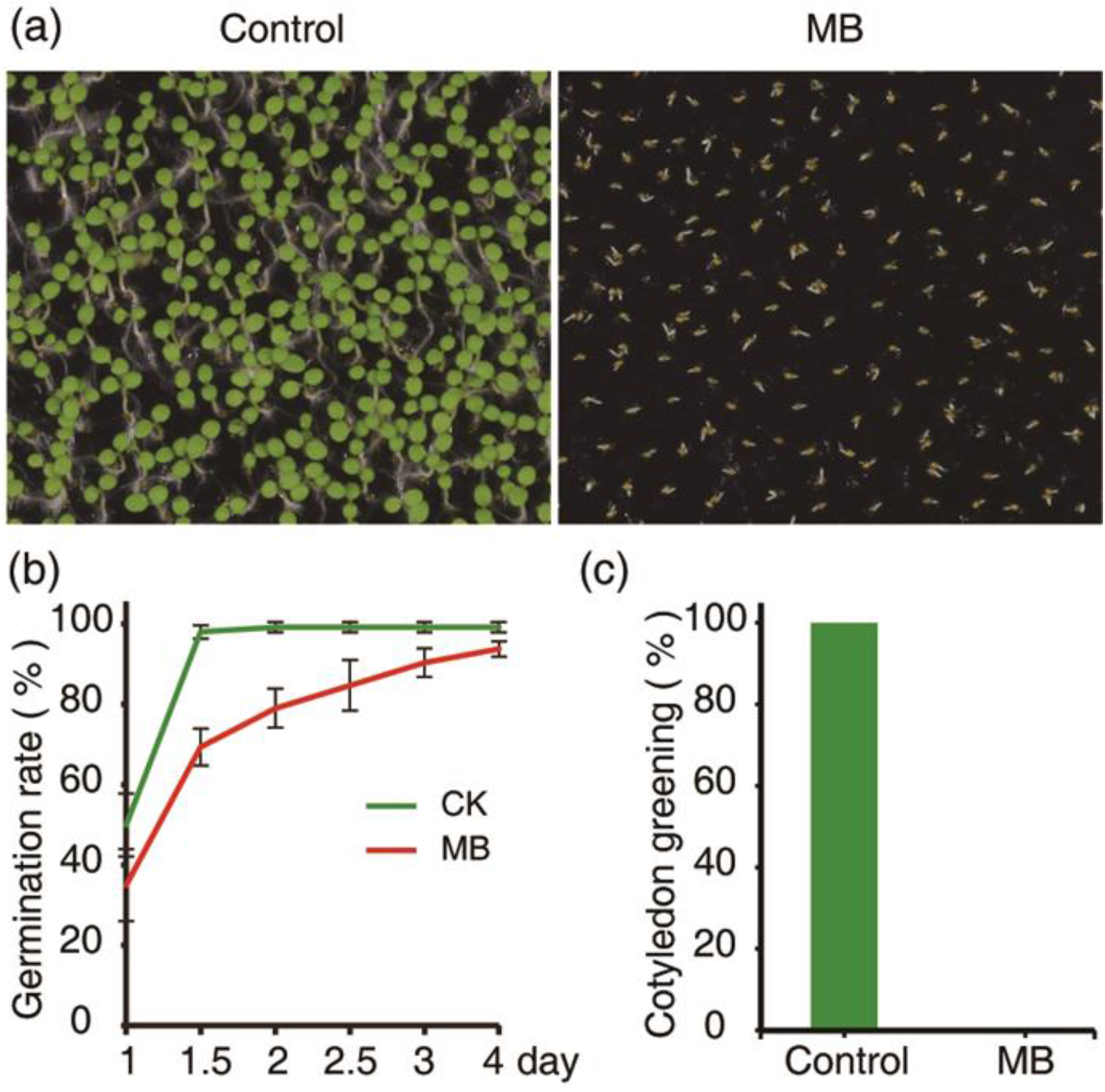
MB inhibits seed germination and seedling establishment of Arabidopsis. Seeds of wild-type Arabidopsis ecotype Columbia were plated on half strength MS medium with or without 4 μM MB, and seed germination and cotyledon greening were examined at the indicated time points. (a) Phenotypes of seedlings 4 d after MB treatment. (b) Percent germination shown at 12 h intervals. (c) Percent cotyledon greening 4 d after MB treatment. Values are mean ± SD from five biological replicates.

### Role for ABA in MB-mediated inhibition of germination and early seedling establishment

Several plant hormones are involved in controlling seed germination, and abscisic acid (ABA) and gibberellin acid (GA) play particularly prominent roles (Shu *et al.*, 2016). The antagonistic interaction between ABA and GA in this regard has been well documented, with ABA involved in the maintenance of seed dormancy and inhibition of germination, while GA breaks seed dormancy and induces cellular processes required during germination (Tuan *et al.*, 2018). To examine the potential roles played by ABA and GA in the inhibition of seed germination by MB, we first analyzed transcript levels of selected genes involved in ABA and GA biosynthesis and signaling in Arabidopsis seedlings germinated in the presence and absence of 4 μM MB (Fig. 2). The transcript levels for *NCED3, NCED6 and NCED9*, which encode rate-limiting enzymes within the ABA biosynthetic pathway, were dramatically increased in MB-treated seedlings relative to untreated control seedlings (Fig. 2a). The ABA-responsive transcription factors *ABI3*, *ABI4* and *ABI5* positively regulate ABA signaling during seed development and germination. Loss of function of *ABI3*, *ABI4* or *ABI5* releases the inhibitory effect of ABA on seed germination (Finkelstein, 2013). *EM1* and *EM6* encode proteins associated with embryogenesis, and can be reactivated by exogenous ABA application during seed germination (Hu *et al.*, 2019). *RD29A* is a typical ABA-responsive marker gene (Nakashima *et al.*, 2006). The levels of *ABI3*, *ABI4*, *ABI5*, *EM1*, *EM6* and *RD29A* transcripts were also markedly increased in MB-treated seedlings relative to controls (Fig. 2c-d). In the case of *ABI4*, the observed transcript level increase was more than one thousand-fold (Fig. 2c), indicating that *ABI4* could play a significant role in the MB-associated inhibition of seed germination.

**Figure 2.**
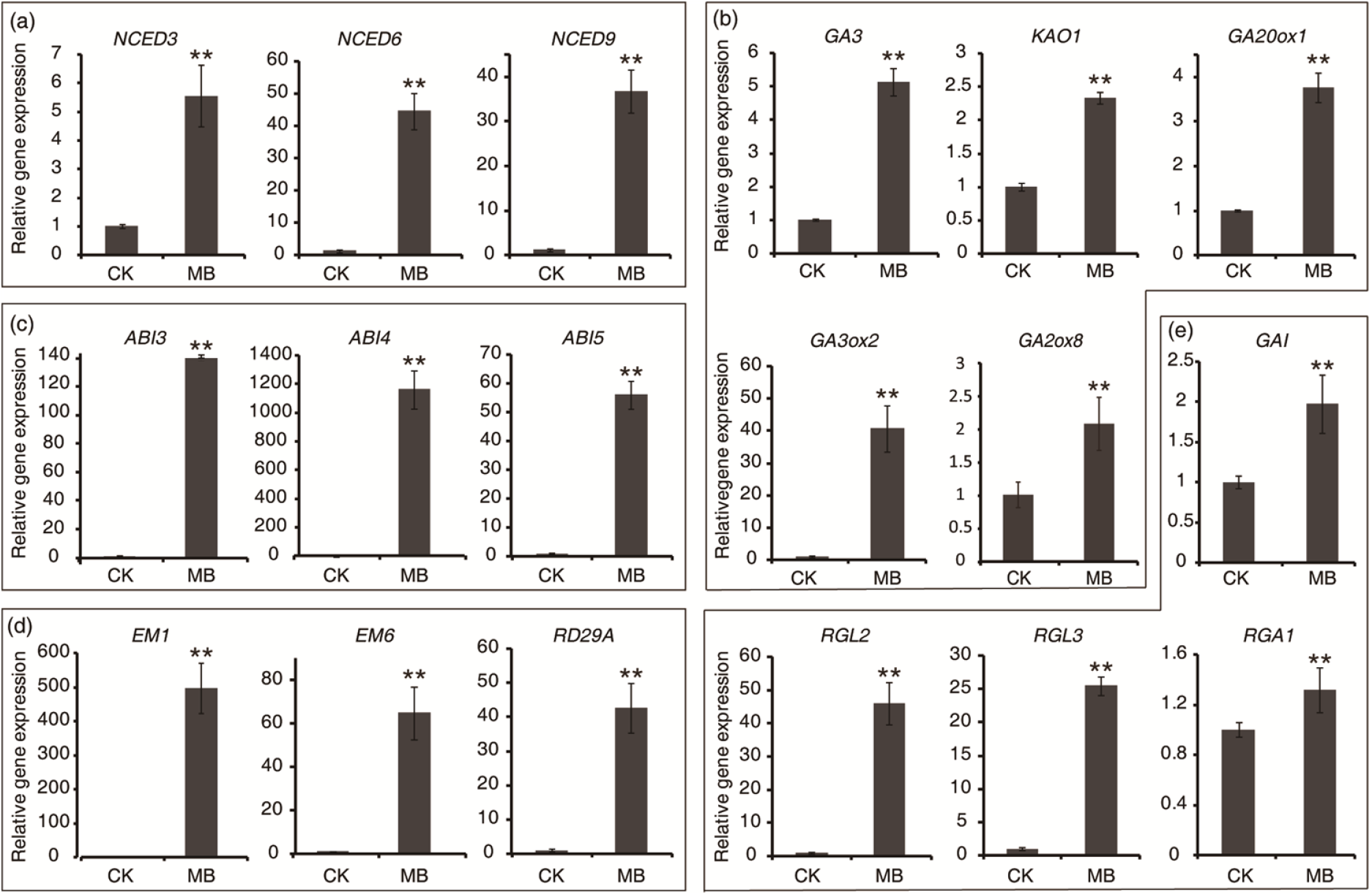
Effect of Momilactone B (MB) exposure on ABA and GA pathway-related transcript levels in Arabidopsis. Seeds of wild-type Arabidopsis were plated on half strength MS medium in the presence or absence of 4 μM MB. 4-day-old seedlings were harvested. Relative transcript levels of representative genes involved in ABA biosynthesis (**a**), GA biosynthesis (**b**), ABA signaling (**c**), ABA response (**d**), and GA signaling (**e**) were monitored by real-time qRT-PCR. The relative gene expression values were normalized to that of the internal control ACTIN2, and then calculated by comparing the value with that of control treatments. Values are mean ± SD from three independent biological replicates. Asterisks indicate significant differences between MB-treated and control (CK) plants (**, *P* < 0.01; Student’s t-test).

To further examine the potential involvement of *ABI4* in the inhibitory effects of MB, Arabidopsis *abi4* mutant seedlings were germinated in the presence of 0, 2, or 4 μM MB and compared with identically-treated wild-type seedlings (Fig. 3). The observed germination and cotyledon greening frequencies of *abi4* mutant plants were similar to WT(At) plants on medium lacking MB. In both the 2 μM and 4 μM MB treatments, the % germination and cotyledon greening of *abi4* mutant seedlings were significantly higher than those of WT plants, indicating that the *abi4* seedlings were more resistant to the inhibitory effects of MB (Fig. 3a-c).

**Figure 3.**
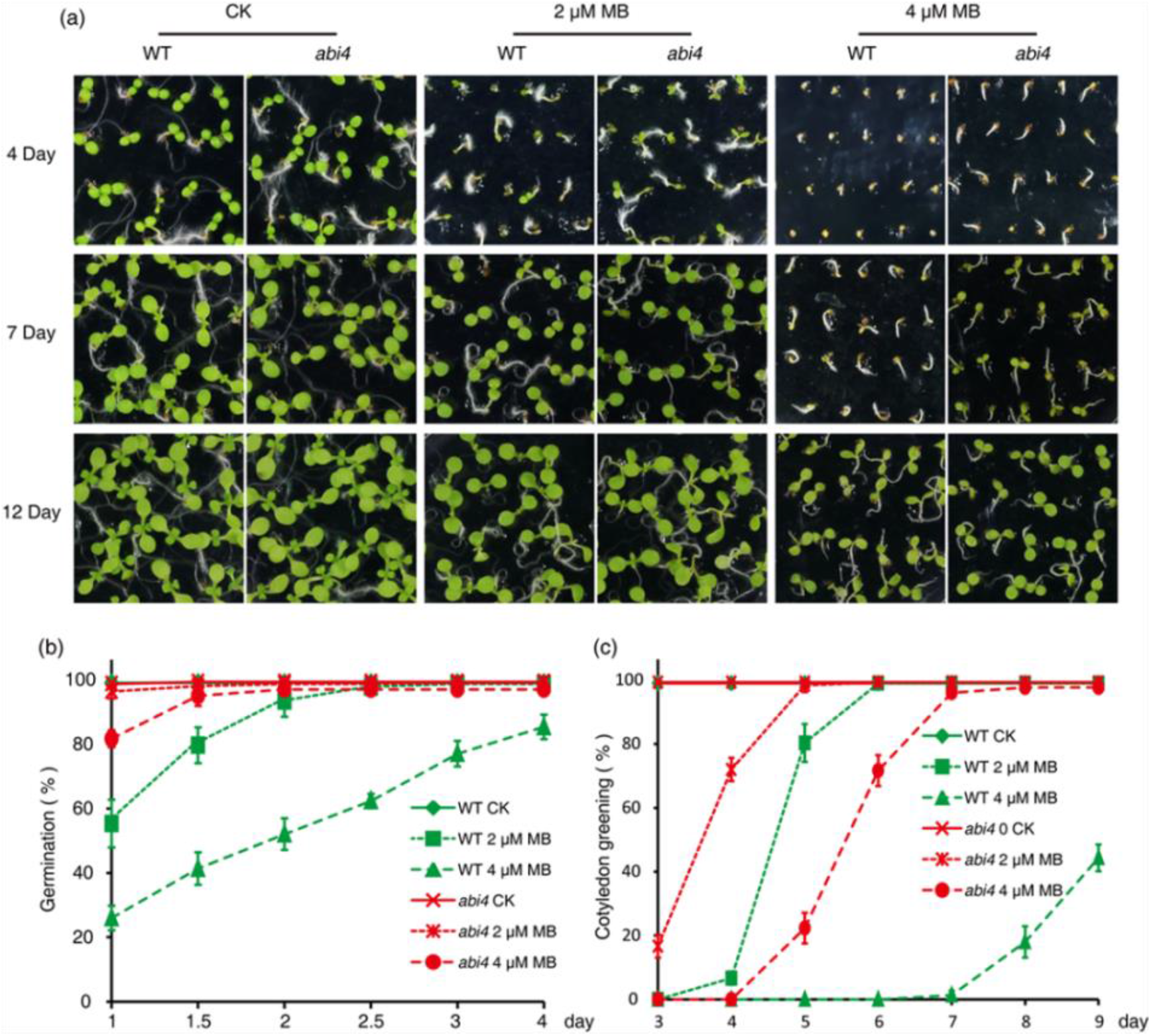
Deficiency in ABI4 increases tolerance of Arabidopsis to momilactone B (MB). Seeds of WT(At) and ABA deficient mutant *abi4* were plated on half strength MS medium containing either 0, 2 or 4 μM MB. Percentage of seed germination and cotyledon greening were determined at the indicated time points. (**a**) Phenotypes of WT(At) and *abi4* seedlings are shown in response to different control or momilactone B treatments. Percent seed germination (**b**) and cotyledon greening (**c**) are shown at different time points for control and momilactone B treatments. Values are mean ± SD from five biological replicates.

Transcript levels of selected genes involved in GA biosynthesis and signaling were also analyzed in seedlings germinated in the presence and absence of 4 μM MB (Fig. 2). DELLA proteins are key negative regulators of the GA-GID1-DELLA signaling pathway and appear to repress all GA-promoted processes, including seed germination and seedling establishment (Sun, 2008). There are five genes encoding DELLA proteins in Arabidopsis, *RGA1*, *GAI*, *RGL1*, *RGL2* and *RGL3*. Our results revealed that transcript levels of the represented DELLA genes, *RGA1*, *GAI*, *RGL2* and *RGL3* were highly up-regulated in MB-treated seedlings relative to untreated controls (Fig. 2e). Interestingly, the transcript levels of representative GA biosynthetic genes, which included *GA3*, *KAO1*, *GA20ox1*, *GA3ox1* and *GA2ox8*, were also increased in MB-treated seedlings (Fig. 2b), presumably via a feedback loop regulating this pathway (Middleton *et al.*, 2012). Given that the transcript levels of *DELLAs* were up-regulated by MB treatment, we performed additional experiments to determine whether co-application of exogenous GA could reverse the inhibitory effects on seed germination caused by MB exposure. WT seeds were germinated on half strength MS medium supplemented with either 2 μM MB, 40 μM GA or 2 μM MB plus 40 μM GA, as well as untreated controls (Fig. 4). At 1.5 day post-planting however, GA addition did not appear to reverse the inhibitory effects of MB, and in fact GA addition exacerbated the inhibition, leading to further decrease in % germination (25.3% for MB plus GA compared with 60.5% for MB treatment alone; Fig. 4a-b). Similar results were obtained for cotyledon greening when GA was co-applied with MB (Fig. 4a, c). At nine days post-planting, the % cotyledon greening was 4.4% in GA plus MB treated seedlings, whereas the observed frequency of greening was 96.4% for seedlings grown in the presence of MB alone. These results suggest that MB inhibits seed germination and seedling establishment, at least in part, by disrupting ABA biosynthesis and/or signaling, however the involvement of GA is unclear.

**Figure 4.**
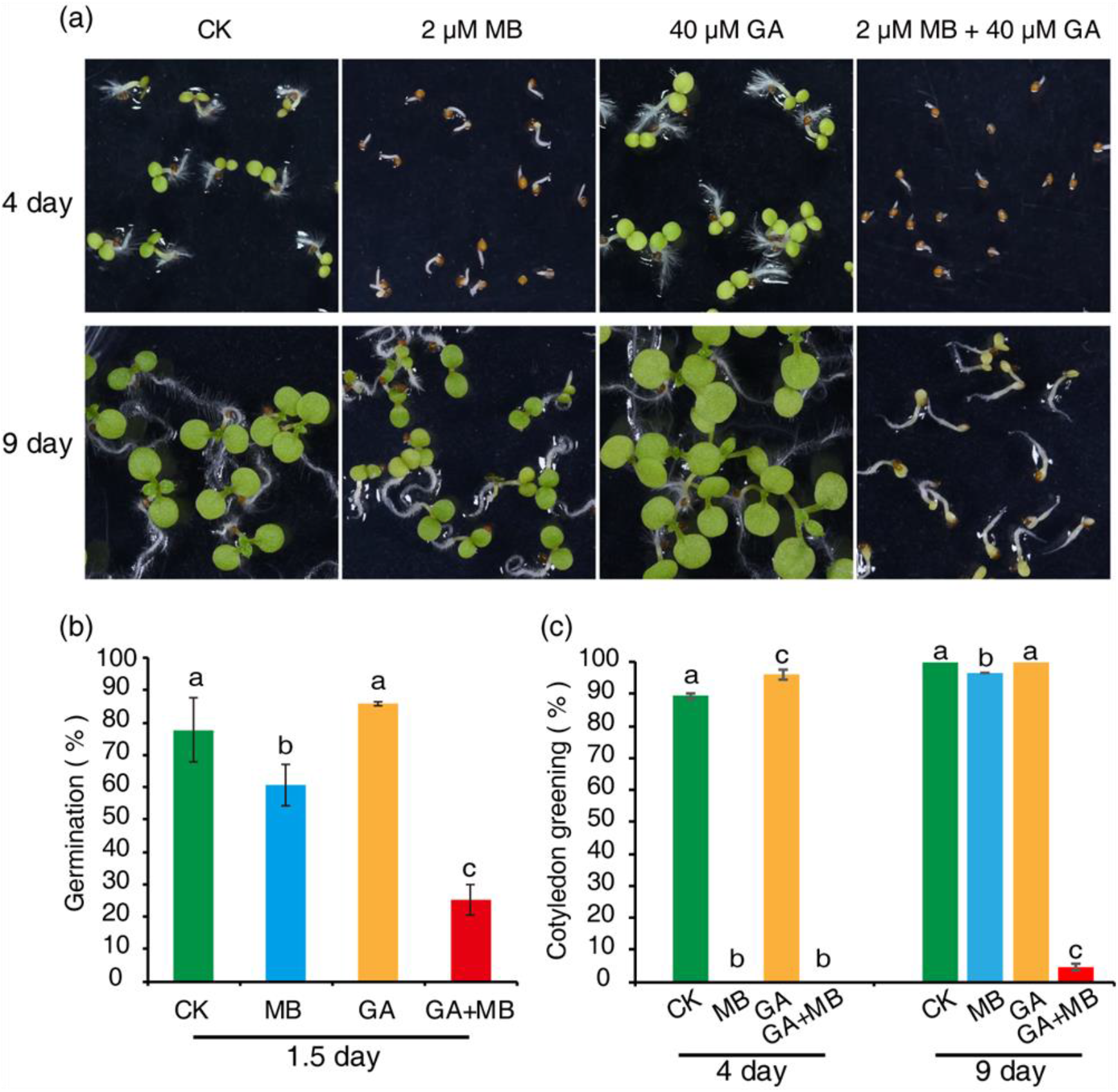
Exogenous GA is unable to reverse the inhibitory effects of momilactone B (MB) on Arabidopsis seed germination. Seeds of wild-type Arabidopsis ecotype Columbia were germinated on half strength MS medium supplemented with 2 μM MB, 40 μM GA and their combination. (**a**) Phenotypes of WT(At) under various treatments; (**b**) percent seed germination, (**c**) percent cotyledon greening. Values are mean ± SD from five biological replicates. Letters above bars indicate significant differences among groups (*P*<0.05, Student–Newman–Keuls test).

### Receiver seedling root systems are sensitive targets for inhibition by MB

In rice, *syn*-copalyl diphospate (*syn*-CDP) synthase (OsCPS4), is an essential enzyme in momilactone biosynthesis which catalyzes the formation of *syn*-copalyl diphosphate (*syn*-CPP) from the general diterpenoid precursor (E,E,E)-geranylgeranyl diphosphate (GGPP) (Xu *et al.*, 2012). To examine the role played by momilactones in the allelopathic potential of rice, an *oscps4* mutant was first created using the CRISPR-Cas9 genome editing method (Xie *et al.*, 2017). The resulting *oscps4* mutant contained a frameshift within the *OsCPS4* coding region, resulting from a single nucleotide deletion (Fig. S1a). *In vitro* allelopathic activity assays were then performed for both wild-type and *oscps4* rice seedling donors against *Echinochloa crus-galli* (barnyard grass) seedlings, a noxious weed frequently found in cultivated rice fields (Khanh *et al.*, 2007). As shown in Fig. 5a-c, deficiency in OsCPS4 reduced the allelopathic activity of rice against barnyard grass seedlings, leading to increased root lengths in the co-cultivated receiver plants, in comparison with the receiver plants co-cultivated with wild-type rice. Consistent with these observations, exogenously provided MB significantly inhibited the growth of barnyard grass seedling root systems at 1, 2 and 4 μM concentrations, whereas seedling shoot system development was relatively unaffected by these treatments (Fig. 5d-e). In addition, comparable results were obtained for pilot assays utilizing *Lactuca sativa* (lettuce) seedlings, a receiver species frequently employed to assess allelopathic activity (Fig. S1b-c). Collectively, the results from the allelopathic activity assay experiments both confirm the significant role played by momilactones in rice allelopathy.

**Figure 5.**
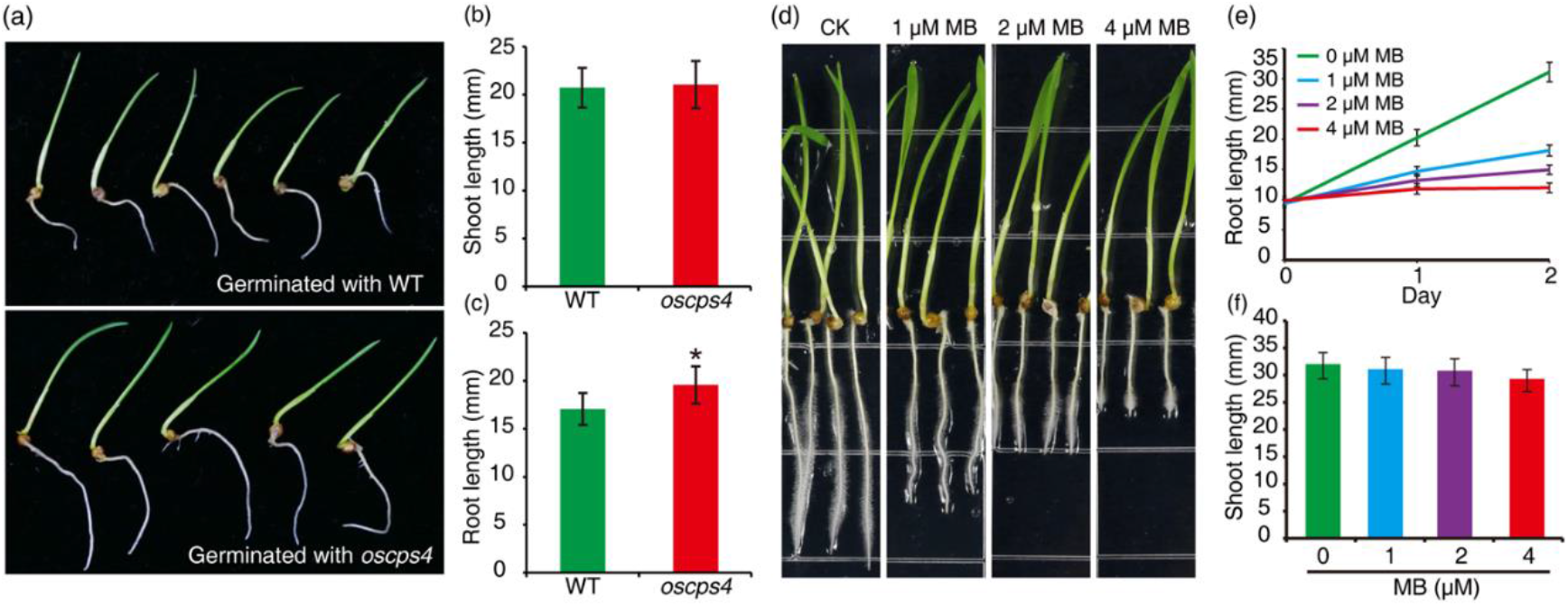
Momilactone B (MB) inhibits root development of barnyard grass. Phenotypes (**a**), shoot (**b**), and root (**c**) lengths of barnyard grass seedlings co-cultured with WT(Os) and *oscps4* knockout rice plants. Phenotypes (**d**), root (**e**), and shoot (**f**) lengths of barnyard grass seedlings treated with MB for 2 d. For barnyard grass values are mean ± SD from 25 seedlings. For MB treatment s, values are mean ± SD from 15 seedlings. Asterisks indicate significant differences (*, P < 0.05; Student’s t-test).

### MB affects Arabidopsis root patterning and primary root meristem maintenance

To further investigate the effect of MB on Arabidopsis root development, 5-day-old seedlings of WT(At) were treated with either 0, 2 or 4 μM of MB, and examined by both confocal and bright field microscopy (Fig. 6). Consistent with the observed effects of MB on barnyard grass (Fig. 5d-e), the growth of Arabidopsis primary roots were significantly inhibited by MB exposure (Fig. 6a, e). Moreover, MB exposure led to a significant reduction in the number of lateral roots produced relative to untreated seedlings, as well as a dramatic reduction in root hair formation (Fig. 6a, b, e, f).

**Figure 6.**
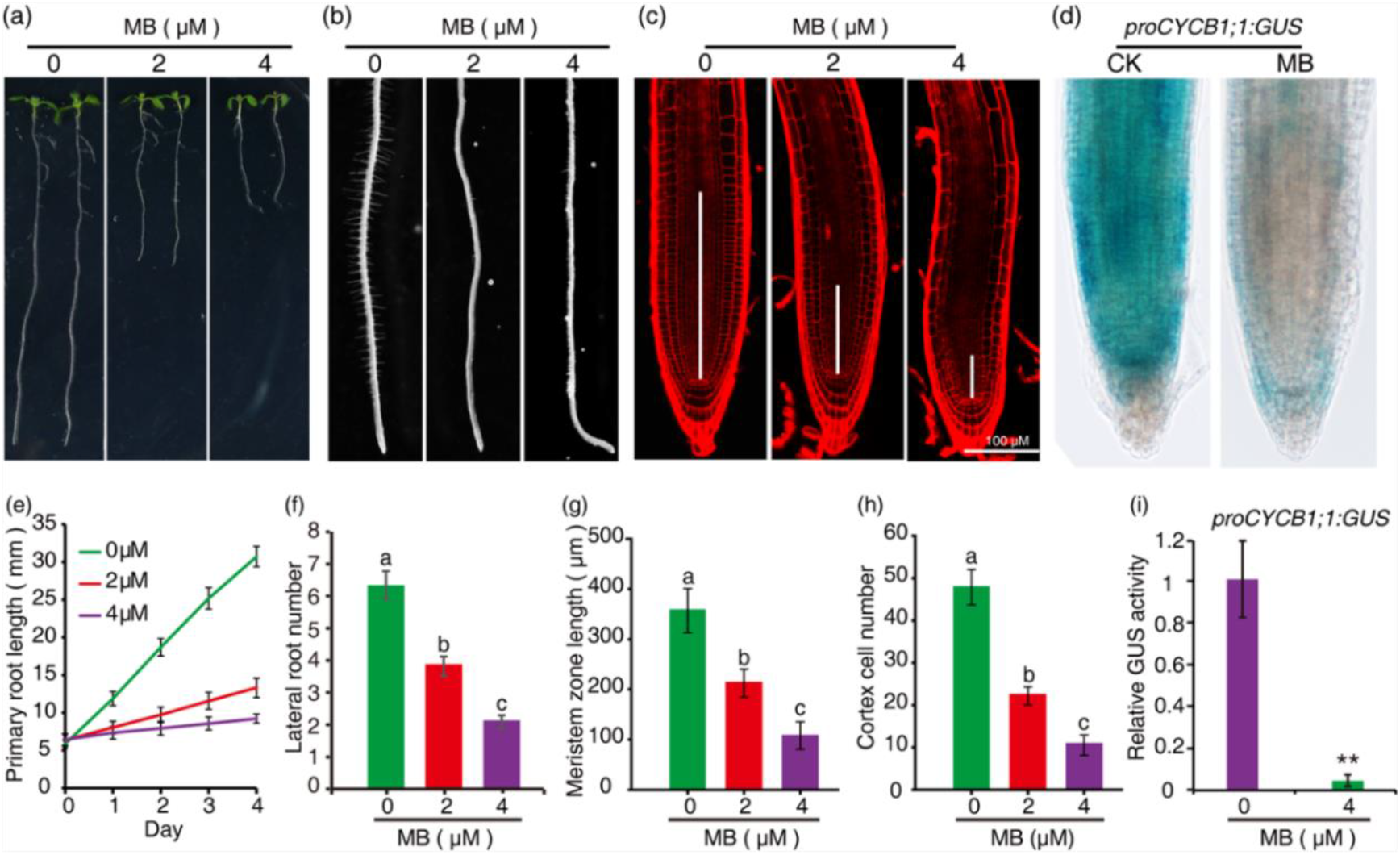
MB-mediated inhibition of Arabidopsis root development. (a) Phenotypes of wild-type Arabidopsis ecotype Columbia WT(At) seedlings treated with various concentration of MB. (b) Root morphology of seedlings described in (a). (c) Root meristem of seedlings described in (a). (d) Histochemical analysis of GUS activity in roots of *proCYCB1;1:GUS* plants in response to 4 μM MB. Statistical analyses of primary root length (e), lateral root number (f), root meristem zone length (g) and root cortical cell numbers (h) of seedlings described in (a). (i) Statistical analysis of GUS activity in roots shown in (d). For quantitative analyses of the root phenotypes, values represent mean ± SD f rom at least 15 seedlings. For GUS activity analysis, at least 8 seedlings were tested, and the values representing mean ± SD are shown relative to control values. Letters above bars indicate significant differences among treatments (*P* <0.05, Student–Newman–Keuls test). Asterisks indicate significant differences (**, *P* < 0.01; Student’s t-test).

Root growth is precisely controlled by cell division and cell differentiation, with the majority of mitotic activity occurring within the meristematic zone located at the distal end of roots (Petricka *et al.*, 2012). To characterize the growth pattern of MB-exposed roots in greater detail, the meristem size and the number of meristematic cortex cells were measured four days after MB treatment. Interestingly, both the meristem size and the number of cortex cells were reduced in MB-treated seedlings (Fig. 6c, g, h). In the 4 μM MB-treated seedlings, root meristem lengths and cortical cell numbers were reduced by approximately 70% and 78%, respectively, compared with untreated seedlings (Fig. 6c, g, h).

The transition from G2 phase to M phase represents a major checkpoint within the eukaryotic cell division cycle. As the *CYCB1;1* gene is expressed specifically at the G2 to M transition during the cell cycle, *proCYCB1;1:GUS* is widely used as a reporter to monitor mitotic activity in plants (Colon-Carmona *et al.*, 1999). To further examine the effects of MB on the meristematic zone in Arabidopsis roots, 5-day old *proCYCB1;1:GUS* transgenic Arabidopsis seedlings were treated with 0 or 4 μM MB for 24 h, and then histochemical analyses of GUS activity in roots were performed (Fig. 6d, i). These tests revealed a significant reduction in GUS activity in roots of MB-treated seedlings compared with untreated seedlings, indicating that MB exposure leads to a reduction in mitotic activity within the root meristematic zone (Fig. 6d, i).

### MB downregulates the expression of transcription factors involved in maintaining the root stem cell niche identity

In the root meristem, a group of mitotically inactive quiescent center (QC) cells and the surrounding stem cells comprise the stem cell niche. The QC maintains the stem cell activity of the surrounding cells, thus functioning as an organizer of the root stem cell niche (Dinneny & Benfey, 2008). To test whether QC cellular function is impaired in MB-treated roots, the QC-expressed promoter trap line QC25 was employed (Sabatini *et al.*, 2003). Histochemical comparisons of GUS activity in MB-treated and untreated QC25 roots revealed an approximately 85% reduction in GUS expression in MB-treated root tips compared to that in the control plants, indicating an obvious loss of QC identity (Fig. 7a, f). QC cell identity is proposed to be maintained via both the PLETHORA (PLT) and SHORTROOT (SHR)/SCARECROW (SCR) pathways (Petricka *et al.*, 2012). PLT1 and PLT2 encode AP2-domain transcription factors, and the maintenance of the stem cell niche is thought to be PLT dosage-dependent (Galinha *et al.*, 2007). SHR and SCR belong to the GRAS transcription factor family. SHR is expressed in the central vascular tissue and moves into the adjacent cell layers to activate SCR transcription, then together with SCR maintain QC and stem cell identity (Wysocka-Diller *et al.*, 2000; Nakajima *et al.*, 2001; Sabatini *et al.*, 2003). The expression of *proPLT1:PLT1-YFP*, *proPLT2:PLT2-YFP*, *proSHR:SHR-GFP* and *proSCR:GFP* reporters were also monitored in MB-treated and untreated transgenic seedling roots by confocal microscopy (Fig. 7b-j). The results of these analyses revealed that the activities of all four reporter genes were significantly reduced in roots of MB-treated seedlings compared with the activities observed in the untreated controls (Fig. 7b-j), indicating that MB exposure leads to disruptions in both the SHR/SCR and PLT regulatory pathways involved in QC cell identity maintenance.

**Figure 7.**
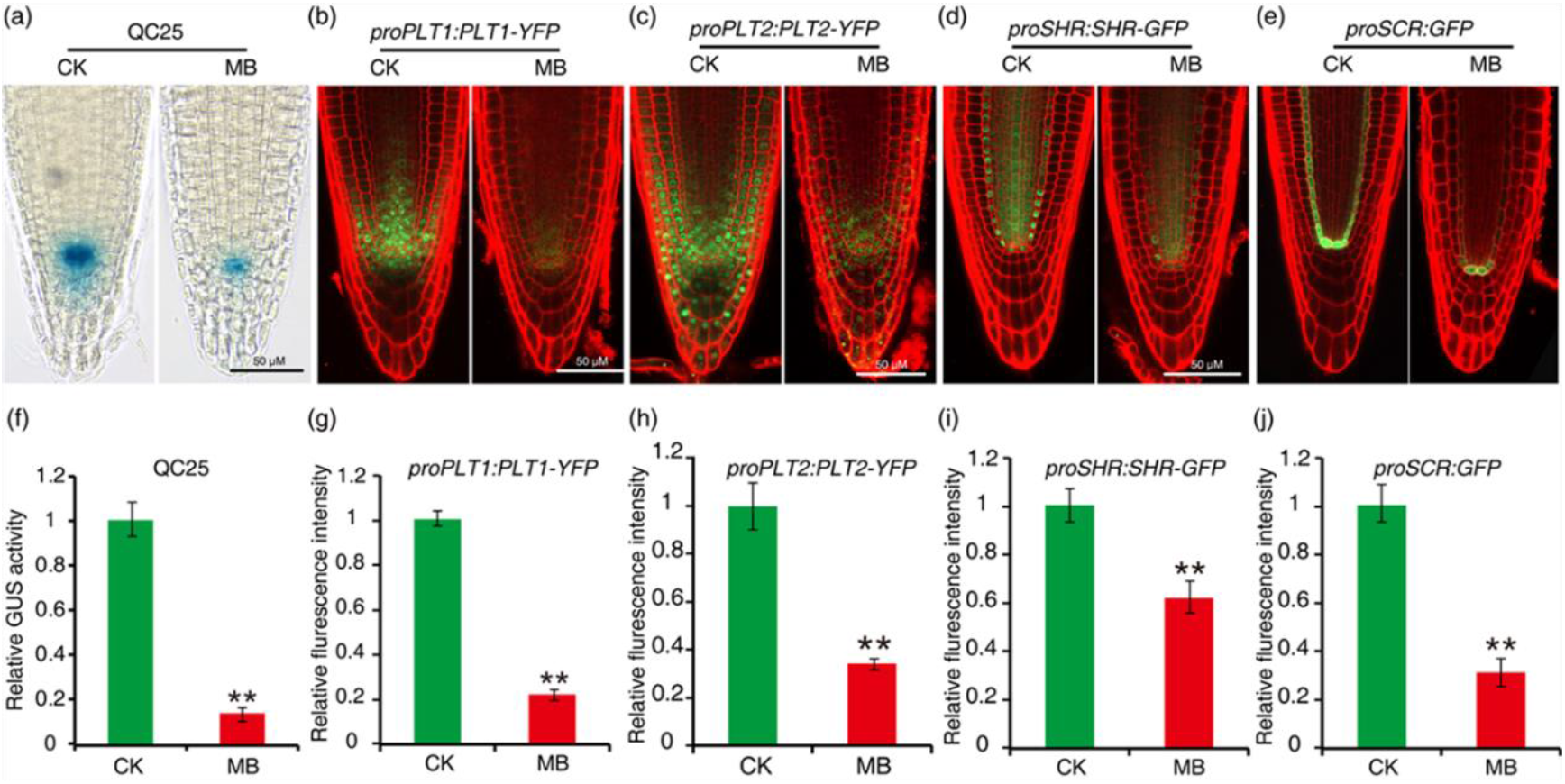
Momilactone B (MB) affects the expression of PLT1/ PLT2 and SCR/SHR in roots. (a) Histochemical analysis of GUS activity in Arabidopsis promoter trap line QC25 in response to 4 μM MB. (b-e) Expression of *proPTL1:PTL1-YFP* (b), *proPTL2:PTL2-YFP* (c), *proSHR:SHR-GFP* (d) and *proSCR: GFP* (e) reporter genes in seedling roots grown for one day in the presence or absence of 4 μM MB. (f) Quantification of GUS activity of roots described in (a). (g-j) Quantification of the florescence intensity of roots described in (b-e). Data represent mean ± SD from at least eight seedlings. Values are shown relative to controls. Asterisks indicate significant differences (**, *P* < 0.01; Student’s t-test).

### MB interferes with auxin biosynthesis and transport in the roots

The expression of PLTs is dependent on endogenous auxin levels, with correct regulation requiring a distribution gradient of PLT protein having a maxima within the stem cell niche (Galinha *et al.*, 2007). The proper distribution gradient of PLTs is essential for meristem maintenance, and correct regulation of cell division and cell differentiation in roots (Galinha *et al.*, 2007). *proDR5:GFP* contains a minimal promoter fused to seven AuxRE repeats driving the expression of *GFP*, and is a reliable marker for monitoring auxin responses and distribution (Friml *et al.*, 2003). DII-VENUS, an auxin sensor, is rapidly degraded in response to auxin and has been used to visualize dynamic changes in cellular auxin distribution (Brunoud *et al.*, 2012). Comparisons between MB-treated and untreated roots using confocal microscopy indicated that expression of the auxin inducible *proDR5:GFP* reporter was reduced by approximately 42% in response to MB exposure (Fig. 8a, f). In contrast, the expression of *DII-VENUS* was increased in MB-treated plants (Fig. 8b, g), indicating that MB exposure is associated with a reduction in auxin content in roots.

**Figure 8.**
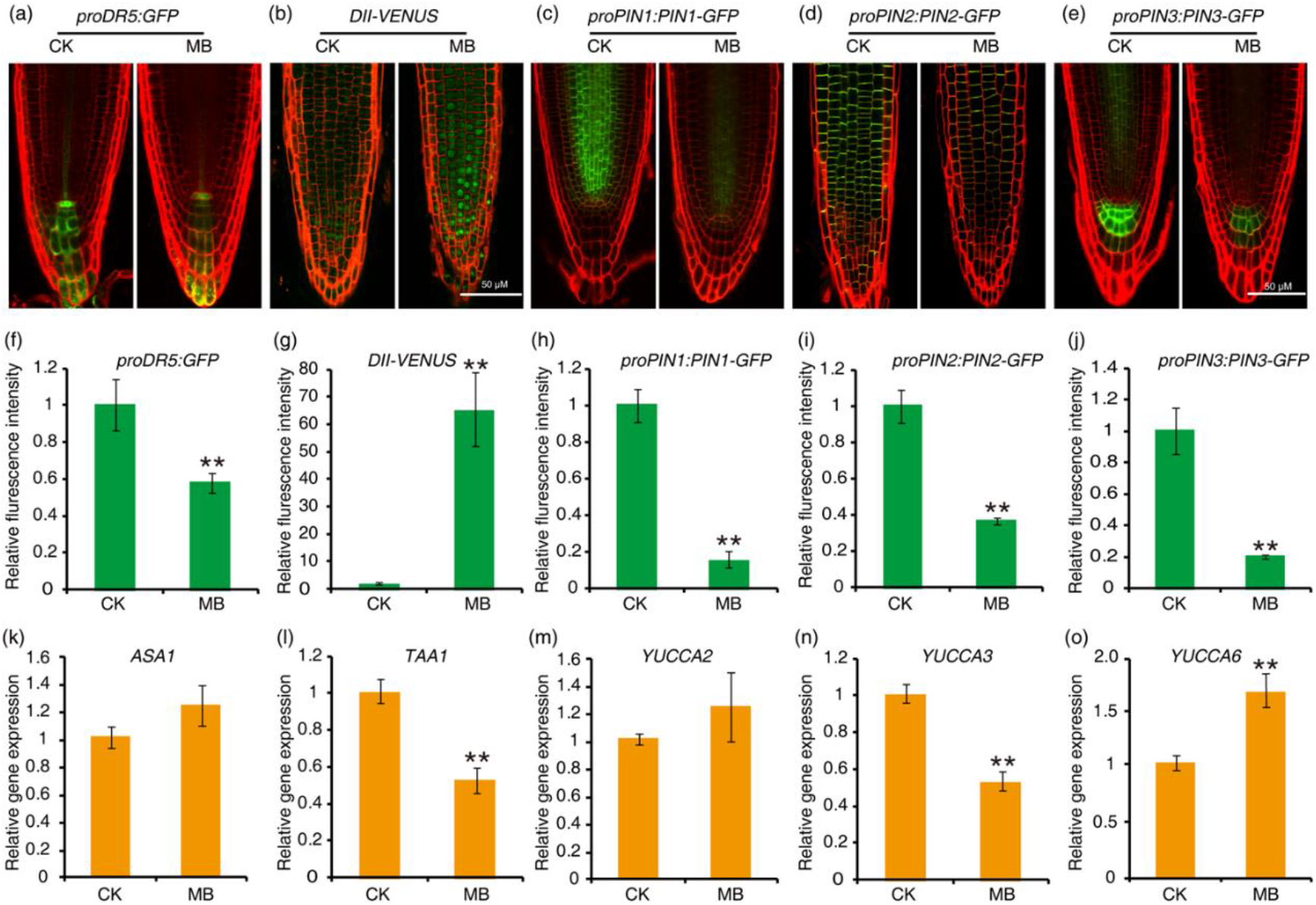
Effects of momilactone B (MB) on auxin pathway in Arabidopsis roots. Expression of *proDR5:GFP* (a), *DII-VENUS* (b), *proPIN1:PIN1-GFP* (c), *proPIN2:PIN2-GFP* (d), and *proPIN3:PIN3-GFP* (e) in root tips grown for 1 d in the presence or absence of 4 μM MB. (f-j) Quantification of florescence intensity in roots described in (a-e). Relative transcript levels for selected auxin biosynthetic pathway genes (k-o) in root tips in response to MB. For quantification of florescence intensity, data represent mean ± SD from at least eight seedlings. Values are shown relative to controls. For qRT-PCR analyses, data represent mean ± SD from three biological replicates. Asterisks indicate significant differences (**, *P* < 0.01; Student’s t-test).

Both local auxin biosynthesis and polar auxin transport are indispensable to root meristem maintenance (Grieneisen *et al.*, 2007; Brumos *et al.*, 2018). The tryptophan-dependent pathway represents the major biosynthetic auxin pathway in Arabidopsis (Mashiguchi *et al.*, 2011). ASA1 (ANTHRANILATE SYNTHASE 1) catalyzes the first reaction of tryptophan biosynthesis, TAA1 (TRYPTOPHAN AMINOTRANSFERASE of ARABIDOPSIS 1) converts tryptophan to indole-3-pyruvic acid (Zhao, 2014), and YUCCA proteins encode flavin monooxygenase which catalyzes the rate-limiting step in tryptophan-dependent auxin biosynthesis – the conversion of indole-3-pyruvic acid to IAA (Mashiguchi *et al.*, 2011). Transcript levels for *ASA1*, *TAA1*, *YUCCA2*, *YUCCA3* and *YUCCA6* were also analyzed in MB-treated and untreated roots by qRT-PCR (Fig. 8k-o). Significant reductions in *TAA1* and *YUCCA3* transcript levels were observed in the MB-treated seedlings relative to the control treatments, whereas *YUCCA6* transcript levels increased (Fig. 8l, n, o). Transcript levels for *ASA1* and *YUCCA2* did not show obvious changes in response to MB exposure.

PIN1, PIN2 and PIN3, together with other PIN proteins, control auxin distribution in Arabidopsis (Friml *et al.*, 2002; Blilou *et al.*, 2005), therefore we also examined the potential effects of MB on the expression of *proPIN1:PIN1-GFP*, *proPIN2:PIN2-GFP* and *proPIN3:PIN3-GFP* reporter genes. Comparisons between MB-treated and untreated roots using confocal microscopy indicated that the expression of the *proPIN1:PIN1-GFP*, *proPIN2:PIN2-GFP* and *proPIN3:PIN3-GFP* reporters were all significantly decreased in roots by MB exposure (Fig. 8c-e, h-j). These results suggest that polar auxin transport may also be disrupted in MB-exposed seedlings through the inhibition of PIN protein expression

Among all PIN proteins, loss of PIN1 function results in the most severe phenotypic effects in Arabidopsis (Blilou et al., 2005). To determine whether the observed potential reduction in PIN protein levels was associated with the proteasome-dependent protein degradation pathway, *pro:PIN1:PIN1-GFP* seedlings were simultaneously treated with the proteasome inhibitor MG132 and MB (Fig. S2). PIN1 levels in roots of MG132 plus MB-treated seedlings did not differ from that of MB-treated seedlings (Fig. S2b). Nitric oxide (NO) has been reported to negatively regulate PIN1 protein levels in a proteasome-independent manner in Arabidopsis (Fernandez-Marcos *et al.*, 2011). We therefore also examined effects of the NO scavenger 2-(4-carboxyphenyl)-4,4,5,5-tetramethylimidazoline-1-oxyl-3-oxide (cPTIO) on PIN1 expression. As was observed for addition of MG132, PIN1 levels in roots of cPTIO plus MB-treated seedlings did not differ from that of MB-treated seedlings (Fig. S2b). Collectively, these results suggest that the observed MB-associated reduction in PIN1 protein levels occurs independently of the proteasome degradation pathway and NO signaling.

### Exogenous auxin partially rescues MB-induced growth inhibition

Given the present results that the content of auxin was reduced in the MB-treated root tips, it is reasonable to speculate that MB-induced growth inhibition could be reversed by exogenous application of auxin. Arabidopsis seedlings were treated with MB plus various concentrations of the synthetic auxin naphthaleneacetic acid (NAA; Fig. 9). Co-application of 10 nM NAA plus 2 μM MB showed a slight, but significant increase in primary root length compared with MB treatment alone (Fig. 9a, c-d). However, root hair production in seedlings grown under these conditions was severely reduced relative to untreated seedlings, and did not differ significantly from that observed in seedlings treated with MB alone (Fig. 9b). Surprisingly, in the 100 nM NAA plus MB treatment, root hair production was fully rescued by the addition of NAA from MB inhibition (Fig. 9b). These results reveal that exogenously-supplied auxin can partially rescue the inhibitory effects of MB on Arabidopsis root development.

**Figure 9.**
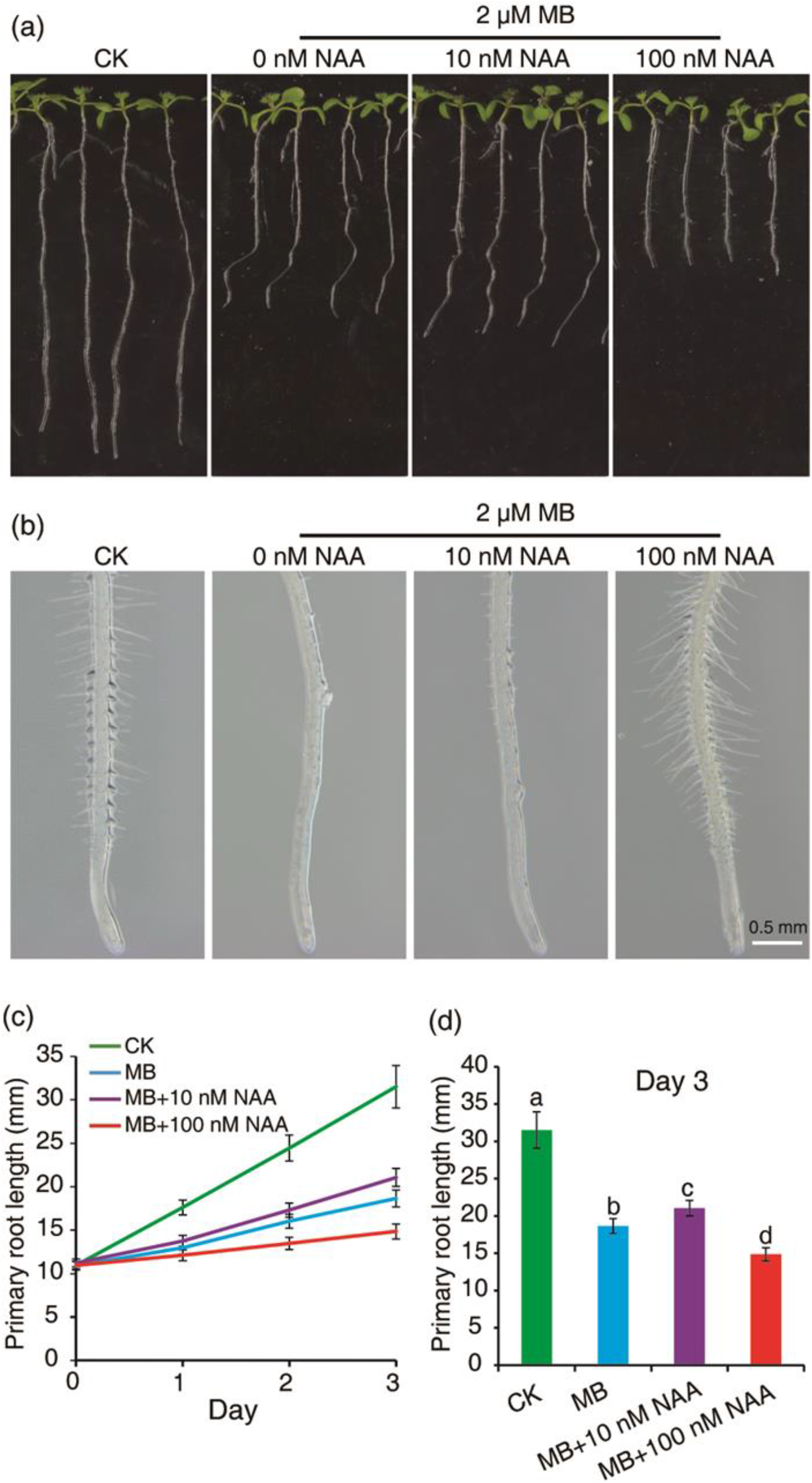
NAA partially rescues the inhibitory effects of momilactone B (MB) on root growth. (a) Phenotypes of wild-type Arabidopsis seedlings treated with various concentrations of NAA and 2 μM MB for 3 days; (b) Phenotypes of the roots described in (a) at higher magnification; (c-d) Statistical analyses of the primary root lengths described in (a). Values represent mean ± SD from at least 15 seedlings. Letters above bars indicate significant differences among treatments (*P*<0.05, Student–Newman–Keuls test).

## Discussion

The identification and characterization of allelochemicals is central to our understanding of plant-plant allelopathy, and the ecophysiological roles played by these compounds (Einhellig & Souza, 1992). Some of the most extensively studied examples include the phenolic lipid sorgoleone, benzoxazinoids such as 4-dihydroxy-2H-1,4-benzoxazin-3(4H)-one (DIBOA), and the diterpene momilactones. Sorgoleone is released from the root hairs of members of the genus *Sorghum*, and inhibits seedling growth in numerous weed species at 10 μM concentrations, causing reductions in biomass accumulation in both the root system as well as aerial plant parts (Einhellig & Souza, 1992). In maize plants, the compound 2-Amino-3H-phenoxazin-3-one (APO) is a degradation product derived from the root system-exuded allelochemical DIBOA, and displays strong inhibitory effects on seed germination and the growth of root and shoot systems of receiver plants *in vitro* at concentrations above approximately 100 μM (Macias *et al.*, 2005; Venturelli *et al.*, 2015). Numerous studies have identified MB as the most active allelochemical secreted from the roots of rice plants (Kato-Noguchi *et al.*, 2002; Kato-Noguchi *et al.*, 2010; Xu *et al.*, 2012). In the present work, we found that MB displayed inhibitory effects on seed germination and primary root growth in Arabidopsis at concentrations as low as 2 μM when tested *in vitro* (Fig. 5, 6). Moreover, MB inhibited lateral root emergence, root hair production, cotyledon greening, and also delayed the initiation of true leaves (Fig. 1 & Fig. 6). Experiments utilizing a CRISPR-Cas9 genome editing-derived rice *OsCPS4* knockout line (deficient in momilactone biosynthesis) also exhibited a significant reduction in allelopathy against barnyard grass, relative to wild-type rice plants (Fig. 5). As the phytotoxic activity of MB is in some cases much greater than that reported for other allelochemicals (Kato-Noguchi & Peters, 2013; Kong *et al.*, 2019), MB could also potentially serve as a natural products-based herbicide.

The inhibition of germination of receiver plants is frequently found to be a target in plant-plant allelopathic interactions. Hormones, especially GA and ABA, play central roles in the regulation of seed germination and have also been shown to be involved in the inhibition of seed germination by allelochemicals (Tuan *et al.*, 2018). For example, the accumulation of ABA in tomato seeds was observed after treatment with the aqueous leachate of the allelopathic weed *Sicyos deppei* G. Don (Cucurbitaceae), leading to delayed germination (Lara-Nunez *et al.*, 2009). Myrigalone A, an allelochemical isolated from fruit leachates of *Myrica gale* L. (sweet gale), inhibits *Lepidium sativum* seed germination by interference with GA biosynthesis and increasing apoplastic reactive oxygen species (ROS) production (Oracz *et al.*, 2012). Recently coumarin, which has been reported to play a role in allelopathic interactions, was also shown to inhibit rice seed germination by suppressing ABA catabolism and promoting ROS accumulation (Chen *et al.*, 2019). Additionally, seed storage protein mobilization, a process regulated by ABA, has been shown to be impaired in response to MB treatment in germinating seeds (Kato-Noguchi *et al.*, 2013), suggesting that ABA may be involved in the MB-mediated inhibition of seed germination. Despite these reports however, relatively little direct genetic evidence exists concerning the potential roles played by phytohormones in allelopathic interactions. In the present study we found that MB treatment markedly increased the expression of the ABA biosynthetic genes *NCED3*, *NCED6* and *NCED9*, as well as the ABA signaling pathway genes *ABI3*, *ABI5*, *EM1*, *EM6*, *RD29A* and *ABI4* (Fig. 2). Genetic analyses showed that loss of *ABI4* function significantly reduced the inhibition of seed germination caused by MB exposure (Fig. 3). Although the expressions of several *DELLA* genes were increased in MB-treated Arabidopsis seedlings, exogenously-provided GA did not rescue the inhibitory effects of MB on seed germination (Fig. 4). In fact, GA addition appeared to exacerbate the inhibitory effects of MB on seed germination, via an unknown mechanism. Thus, the present results clearly support a role for ABA biosynthesis and/or signaling in the mechanism underlying MB-mediated germination inhibition, whereas a potential role for GA is less evident.

For some plant-plant allelopathic interactions, the inhibition of receiver seedling root system growth may occur in the absence of any obvious inhibitory effects on germination. In fact, a number of putative allelochemicals exhibit this type of activity, including sorgoleone, coumarin, benzoic acid and cyanamide (Einhellig & Souza, 1992; Soltys *et al.*, 2012; Lupini *et al.*, 2014; Zhang *et al.*, 2018). For example, sorgoleone does not affect the germination of the weed *Eragrostis tef*, but does suppress root and shoot system growth in *Eragrostis* seedlings (Einhellig & Souza, 1992). The inhibitory effect of sorgoleone on root growth involves a reduction in the activity of root H+-ATPase as well as delayed cell division (Hejli & Koster, 2004). Coumarin is also a putative allelochemical that is widely-distributed throughout the plant kingdom. Coumarin has been shown to inhibit primary root elongation and stimulate lateral root formation (Lupini *et al.*, 2014). Mutation of the root-specific auxin influx transporter AUX1 rescues root growth inhibition by coumarin, indicating that coumarin might act via interference with polar auxin transport (Lupini *et al.*, 2014). Benzoic acid is another putative allelochemical found within the root exudates of numerous plant species. Exposure to this compound up-regulates the expression of auxin biosynthetic genes as well as the auxin polar transporter genes *AUX1* and *PIN2* in roots, resulting in increased auxin levels which may be responsible for the observed benzoate-mediated inhibition of primary root growth (Zhang *et al.*, 2018). Thus, auxin biosynthesis and/or signaling could represent processes which are frequent targets for allelochemicals.

Consistent with prior studies performed by (Kato-Noguchi *et al.*, 2012), we found that MB strongly inhibited primary root growth, and in the present work we also show this inhibition is associated with a substantial reduction in root meristem activity (Fig. 7). Stem cell niche activity within the root meristem is specified and maintained by two parallel pathways: the PLT pathway and the SHR/SCR pathway (Cederholm *et al.*, 2012). Auxin is a key regulator in stem cell positioning, and forms a gradient distribution pattern within the root meristem (Cederholm *et al.*, 2012). The asymmetric distribution of auxin is generated and maintained by the auxin transporters (Blilou *et al.*, 2005). In response to auxin within the root meristem, PLT proteins display a gradient which guides the progression of cells from stem cell state, to transit-amplifying cell state, and finally to differentiation (Galinha *et al.*, 2007). PLT proteins regulate the expression of PIN genes to stabilize the auxin gradient in the root meristem (Galinha *et al.*, 2007). The SHR/SCR pathway controls the radial positioning of the quiescent center and is auxin-independent (Wysocka-Diller *et al.*, 2000; Nakajima *et al.*, 2001). In this study, we found that MB treatment downregulated the expressions of *QC25, PLT1/PLT2* and *SHR/SCR* (Fig. 7). Auxin content in root tips was monitored by the auxin sensor DII-VENUS, and the auxin responsive reporter *proDR5:GFP* (Brunoud *et al.*, 2012). Both assays indicated that MB exposure led to reduced auxin content in root tips (Fig. 8a-b, f-g), which is essential for root meristem maintenance (Brumos *et al.*, 2018). qRT-PCR analyses of representative auxin biosynthetic pathway genes, particularly *TAA1* and *YUCCA3*, indicated that auxin biosynthesis is inhibited by MB (Fig. 8k-o). The levels of PIN proteins PIN1, PIN2 and PIN3 were dramatically decreased after MB treatment (Fig. 8c-e, h-j). Application of the proteasome inhibitor MG132 or the nitrous oxide scavenger cPTIO could not reverse the observed reduction of PIN1 protein in MB-treated root tips, indicating that this occurred independently of the proteasome pathway and nitrous oxide. In addition, exogenous auxin partially restored root growth in the presence of MB, and fully restored the production of root hairs (Fig. 9). Taken together, these results suggest that the inhibitory effects of MB on root growth may also involve the disruption of auxin biosynthesis and PIN-mediated polar auxin transport.

In summary, this study provides significant insights into the mechanisms underlying the inhibition of germination and early seedling establishment by momilactone B. MB-mediated interference of plant growth may involve multiple cellular targets, including the phytohormones ABA and auxin and their respective biosynthetic and signaling pathways. Our results suggest that MB-mediated inhibition of seed germination may occur via induction of the ABA signaling pathway and ABA biosynthesis, and the inhibition of root growth by MB could be due, at least in part, to a reduction in auxin production and interference with polar auxin transport, thereby impairing the maintenance of the root apical meristem. Thus, the vital regulatory roles played by various phytohormones during seedling development could render their respective pathways as highly effective targets for allelochemical interference.

## MATERIALS AND METHODS

### Plant materials and growth conditions

The wild-type Arabidopsis ecotype Columbia (WT(At)) and the wild-type genotype rice *Oryza sativa* L. cv. Shishoubaimao (WT(Os)) were used in this study. Some of the plant materials in this study have been described previously: *proCyclinB1;1:GUS* (Colon-Carmona *et al.*, 1999); *QC25:GUS* (Sabatini *et al.*, 2003); *proPLT1:PLT1-YFP* and *proPLT2:PLT2-YFP* (Galinha *et al.*, 2007); *proSHR:SHR:GFP* (Nakajima *et al.*, 2001); *proSCR:GFP* (Wysocka-Diller *et al.*, 2000); *proDR5rev:GFP* (Friml *et al.*, 2003); *DII-VENUS* (Brunoud *et al.*, 2012); *pro:PIN1:PIN1-GFP* (Benkova *et al.*, 2003); *pro:PIN2:PIN2-GFP* (Blilou et al., 2005); *pro:PIN3:PIN3-GFP* (Zadnikova *et al.*, 2010)*; abi4* (Shu et al., 2013).

Arabidopsis seeds were surface sterilized and plated on half strength MS medium supplemented with 1% sucrose and 0.7% agar, and then stratified in the dark at 4°C for 2 days before being allowed to germinate at 22°C under long-day conditions (16 h of light/8 h of dark). One-week old seedlings were then transferred to soil for growth at 22°C under long-day conditions (16 h of light/8 h of dark) (Wu *et al.*, 2015a). Rice seeds were surface sterilized and germinated on water-soaked filter paper for 3 d in dark at 30°C. The germinated rice seedlings were then planted in rice paddy fields.

### Isolation of MB

Momilactone B was isolated from rice hulls in our laboratory as described by Gu et al. (2019). Its structure was confirmed by using 1D and 2D NMR in combination with ESI-MS and HR-EIMS.

### Seed germination assays

Arabidopsis plants were grown in a green house at 22°C under long-day conditions (16 h of light/8 h of dark). The seeds were harvested and stored in a dry cabinet for 6 months to break seed dormancy (Shu *et al.*, 2013). To analyze the effect of MB on seed germination of Arabidopsis, seeds were surface sterilized for 5 min in 70% (v/v) ethanol, washed three times with sterile water, distributed on half strength MS medium, with 0.7% (w/v) agar, 1% (w/v) sucrose, and MB at the indicated concentrations, and chilled in the dark at 4°C for 2 days before being allowed to germinate at 22°C. Germination frequency was scored based on recognizable radicle protrusion. Seedling morphology was scored based on cotyledon expansion and greening.

### Root growth assays

To examine the effect of MB on root growth of barnyardgrass (*Echinochloa crus-galli* L.) and Arabidopsis, seedlings were grown vertically on half strength MS agar medium for 2 d or 5 d, respectively, and then transferred to fresh medium supplemented with MB at the indicated concentrations. Root and shoot length, as well as lateral root numbers were measured after treatments.

### Plasmid construction and rice transformation

To create the *OsCPS4* (*LOC_Os04g09900*) deficient rice mutant, the CRISPR-Cas9 genome editing method (Xie *et al.*, 2017) was employed to generate the construct *KO*-*OsCPS4*, targeting the site (GTTTGGGCAGCCAGCATCGG) specific for *OsCPS4*. The resulting construct was transformed into *Oryza sativa* L. cv. Shishoubaimao (Toki *et al.*, 2006). The homozygous mutant *oscps4* lacking the CRISPR construct was identified and used for further study.

### Co-culture assay for evaluating rice allelopathy

The assay used for evaluating allelopathic activity was performed as previously described with slight modifications (Xu *et al.*, 2012). Seeds of WT rice and *oscps4* were surface sterilized and germinated on water-soaked filter paper for 3 days. Six uniformly developed seedlings of each species were transplanted to each of 15 plates containing water-soaked filter paper. Ten surface-sterilized receiver plant seeds (lettuce or barnyard grass) in each plate were co-cultured with the seedlings for 6 days.

### Analysis of transcript levels

Arabidopsis seedlings germinated on medium containing 0 or 4 μM MB for 48 h were harvested. Realtime PCR analysis was conducted as described previously with slight modifications (Wu *et al.*, 2015b). Total RNA was extracted using a Plant RNA Kit (R6827, Omega) and then reverse transcribed using PrimeScript™ RT Master Mix (RR036Q, Takara) per the manufacturer’s instructions. The resulting cDNA templates were subject to qRT-PCR analyses using TB Green^®^ Premix Ex Taq™ II (RR820Q, Takara) with a StepOneTM Real-Time PCR System (Applied Biosystems) according to the manufacturer’s instructions. *ACTIN2* (*AT3G18780*) was used as a reference gene. Data presented are means from three biological replicates with SD. Statistical significance was evaluated by Student’s t test. The primers used are listed in Supplemental Table S1.

### Histochemical and microscopy analyses

Histochemical analyses of GUS activities in the roots of CYCB1;1:GUS and QC25 plants were performed according to methods described previously with slight modifications (Jefferson *et al.*, 1987). Whole seedlings were stained in GUS staining solution (1 mg/mL X-glucuronide in 0.1M potassium phosphate, pH 7.2, 0.5 mmol/L ferrocyanide, 0.5 mmol/L ferricyanide, and 0.1% Triton X-100) at 37 °C in the dark for 12 h. After staining, seedlings were cleared and photographed using a Leica DM6 microscope.

For confocal microscopy imaging, Arabidopsis roots were stained with 10 mg/mL propidium iodide (PI) for 5 min, washed once in distilled water, and mounted in water. Samples were captured using a ZEISS LSM800 confocal laser-scanning microscope with the following excitation/emission wavelengths: 561 nm/600 to 655 nm for PI, 488 nm/498 to 552 nm for GFP and 514 nm/530 to 600 nm for YFP respectively (Zheng *et al.*, 2018). Fluorescence intensities were measured using ImageJ (http://imagej.nih.gov/ij/) and statistical significance was evaluated by Student’s t test.

## ACKNOWLEDGMENTS

We thank the ABRC (Arabidopsis Biological Resource Center) and Drs. Ben Scheres, Philip Benfey, Chengwei Yang and Chuanyou Li for sharing their published materials used in this study. We also thank Dr. Yaoguang Liu for providing CRISPR-Cas9 vectors, and Dr. Ying-Tang Lu for very helpful discussions.

## Supplemental Data

**Figure S1**. Generation of momilactone biosynthesis deficient mutant *oscps4* using CRISPR-Cas9. (a) Sequence analysis of the mutated site within the generated *oscps4* knockout line. (b) Phenotypes and (c) length of roots and hypocotyls of lettuce plants co-cultured with wild-type and *oscps4* mutant plants for 6 days. Data represent mean ± SD from 100 seedlings (**, *P* < 0.01; Student’s t-test).

**Figure S2**. Momilactone B (MB) reduces PIN1 protein level in a proteasome pathway and NO-independent manner. Distribution of *pro:PIN1:PIN1-GFP* protein in Arabidopsis seedlings treated with 4 μM MB for 0 to 24 h (a), and in seedlings treated with proteasome inhibitor MG132 (100 μM) or NO scavenger cPTIO (1 mM), with or without 4 μM MB for 24 h (b).

**Table S1**. Primers used in this study.

## Notes

**Funding information:** This research was supported by the National Natural Science Foundation of China (31670414, 31870361), Natural Science Foundation of Fujian Province (2017J01427), Educational Research Program for Young and Middle-aged Teachers of Fujian province (JAT160154), and Guangzhou Science and Technology Innovation Commission (201804010034).

